# *In vitro* and *in vivo* phasor analysis of stoichiometry and pharmacokinetics using short lifetime near-infrared dyes and time-gated imaging

**DOI:** 10.1101/212977

**Authors:** Sez-Jade Chen, Nattawut Sinsuebphon, Alena Rudkouskaya, Margarida Barroso, Xavier Intes, Xavier Michalet

## Abstract

We introduce a simple new approach for time-resolved multiplexed analysis of complex systems using near-infrared (NIR) dyes, applicable to *in vitro* and *in vivo* studies. We show that fast and precise *in vitro* quantification of NIR fluorophores’ short (sub-nanosecond) lifetime and stoichiometry can be done using phasor analysis, a computationally efficient and user-friendly representation of complex fluorescence intensity decays obtained with pulsed laser excitation and time-gated camera imaging. We apply this approach to the study of binding equilibria by Förster resonant energy transfer using two different model systems: primary/secondary antibody binding *in vitro* and ligand/receptor binding in cell cultures. We then extend it to dynamic imaging of the pharmacokinetics of transferrin engagement with the transferrin receptor in live mice, elucidating the kinetics of differential transferrin accumulation in specific organs, straightforwardly differen tiating specific from non-specific binding. Our method, implemented in a freely-available software, has the advantage of time-resolved NIR imaging, including better tissue penetration and background-free imaging, but simplifies and considerably speeds up data processing and interpretation, while remaining quantitative. These advances make this method attractive and of broad applicability for *in vitro* and *in vivo* molecular imaging, and could be extended to applications as diverse as image guided-surgery or optical tomography.

## 1 Introduction

Molecular imaging enables the characterization and quantification of biological processes in live and intact subjects[1–5]. The ability to monitor, with high spatial and temporal resolution, biological processes *in vivo* and longitudinally has been central to improving our knowledge of integrative biology, more sensitive and specific means to characterize diseases, guiding the development of novel therapies and/or rationalizing therapeutic regimen selection, as well as dynamically assessing treatment efficacy. The most important characteristics sought in molecular imaging techniques are the ability to sense minute amounts of molecules as well as being able to simultaneously track various markers in the same tissue. In this regard, Positron Emission Tomography (PET) is the main molecular imaging technique for diagnosis and study in human clinical trials thanks to its high sensitivity, and it is likewise common for preclinical studies. Despite its widespread use, one of the main drawbacks of PET is that the information provided by the pooling of the injected radiotracers can be nonspecific if the tracer is not well-targeted, such as for the clinically-approved marker FDG (fluodeoxyglucose) for tracking of metabolic activity[6]. The sensitivity of PET can also be highly dependent upon the tracer itself and its efficacy of delivery within the body with regards to metabolism and clearance of the tracer[5], and the use of radiotracers limits its potential for longitudinal studies. Lastly, it is difficult to perform PET multiplexing studies (track multiple biomarkers) *in vivo*. Conversely, optical imaging techniques demonstrate in preclinical settings similar levels of sensitivity to PET with the advantage of being able to use a wide array of fluorescence reporters, including genetically-encoded reporters. Though molecular optical imaging has historically been mainly limited to *in vitro* and preclinical work, there has been a surge in the last decades in the development of optical imaging techniques for clinical applications ranging from guided surgery to medical intervention.

Molecular optical imaging methods harness information provided by light reflected from or transmitted through tissues, as well as emitted fluorescence or luminescence from probes (exogenous or endogenous) within the tissue. A wide array of functionalized fluorophore libraries has been developed over the years to permit tracking numerous molecular processes or biomarkers of interests *in vitro* and *in vivo*. However, in the case of *in vivo* applications, optical imaging of intact tissue does not provide cellular or intra-cellular resolution and hence, like PET, is limited to macro- or meso-localization of the fluorescent probe[7, 8]. Thus, for applications involving delivery of molecular probes, such as in targeted drug delivery, only reporting of the probe pooling at the tissue level is possible. This is particularly an issue in oncological applications in which vascular delivery of probes harnesses the enhanced permeability and retention (EPR) effect[9, 10] but does not guarantee intracellular delivery.

In contrast, fluorescence lifetime is a property of a fluorophore (whether exogenous or endogenous), which is mostly independent of concentration but is sensitive to changes to the microenvironment such as pH or ion concentration[11, 12]. In addition, quantitative lifetime imaging is a powerful tool to monitor nanoscale biological interactions via Förster Resonance Energy Transfer (FRET), a phenomenon in which an excited fluorophore (the ‘donor’) transfers energy non-radiatively to a nearby fluorophore (the ‘acceptor’) provided they have sufficient spectral overlap and are 2-10 nm apart[13, 14]. This results in a decrease in donor fluorescence emission intensity and lifetime due to quenching, and an increase in the acceptor intensity. Quantification of this change in the donor lifetime provides direct information on the FRET interaction.

Fluorescence lifetime imaging (FLI) has proven to be a robust tool *in vitro* and *in vivo* for observation of contrast within biological tissues[11, 12, 15–17]. Lifetime imaging is well-established in microscopy and in the visible range of the spectrum, but its application *in vivo* is technically challenging due to significant and diverse background auto-fluorescence and high signal attenuation in tissues. These limitations can be mitigated by using NIR or longer wavelength emitting species, due to reduced scattering and absorption[18], at the expense of much shorter lifetimes. Moreover, to quantitatively retrieve lifetime or the stoichiometry of fluorophores characterized by different lifetimes, model-based analysis is required and usually involves fitting algorithms. Iterative fitting approaches typically used are computationally demanding, and a challenge when applied to short lifetimes in scattering media. All these characteristics constitute obstacles to fast quantitative NIR lifetime imaging, as would be needed in some clinical applications such as guided surgery or to study fast *in vivo* kinetics. For such applications, the availability of fast quantitative tools for lifetime imaging is still lacking. Herein, we report on the use of phasor analysis, a well-known technique for fluorescence lifetime analysis[19–33], to retrieve quantitatively, accurately and in close to real-time, the lifetime and stoichiometry of short lifetime NIR species in small animal whole-body *in vivo* NIR FRET imaging.

NIR FRET is a promising approach to observe cellular interactions *in vivo* and to assess drug delivery efficacy. In this study, we utilize the transferrin/transferrin receptor (Tf/TfR) pair as a model of ligand/receptor binding interaction. Tf is a blood plasma glycoprotein acting as an iron transporter and binding to TfR. TfR is highly expressed in the liver, which is the main organ regulating iron homeostasis, but also in cancerous tissue[34]. This makes Tf an attractive carrier for targeted therapy[35], as binding of two drug-carrying Tf molecules to the receptor dimer results in endocytosis and delivery of the drug into the cell[36, 37]. Here, we use NIR donor and acceptor dyes conjugated to Tf to probe the Tf/TfR receptor-ligand dimerization and internalization process *in vivo*. Due to the close proximity of the TfR binding sites within the dimer, the donor- and acceptor-labeled Tf experience FRET upon binding and internalization, resulting in a reduced (quenched) donor lifetime[38, 39]. The detection of a quenched donor lifetime by FLI within a region of interest thus directly reports on Tf delivery into the cells, as we recently verified *in vivo*[40, 41].

Standard analysis of FLI-FRET data typically consists of fitting the fluorescence lifetime decay in each pixel to a multi-exponential model[42, 43]. This subsequently allows for extraction of FRET parameters such as the FRET efficiency and the fraction of donor quenched due to FRET within the sample. However, fluorescence decay fitting analysis can be time-consuming and fraught with problems when applied to large images with limited signal-to-noise ratio. Alternative methods such as rapid lifetime determination (RLD[44]) and minimal fraction of interacting donor[45] have been developed to circumvent these limitations, but these methods have not been tested with short-lifetime NIR dyes in the presence of broad impulse response function (IRF) such as encountered in *in vivo* measurements[46]. In these situations, an approach such as phasor analysis that does not rely on specific decay models and does not focus on extracting lifetimes, could be particularly useful due to its speed and ease of use as demonstrated next[19–22, 24, 25, 27, 47–50].

Phasor (or polar plot) analysis of fluorescence lifetime images (phFLI) was initially introduced as a simple and graphical way to represent data acquired with frequency-modulated systems[48, 51]. The method can be used as well with fluorescence lifetime data acquired with other modalities, such as scanning confocal time-correlated single photon-counting (TCSPC)[19], wide-field time-gated[20] or wide-field photon-counting[23] detection and others[52]. Briefly, the method consists in transforming the recorded fluorescence decay *F(t)* into a pair of numbers (*g, s*) proportional to the Fourier transform of *F* at a chosen frequency *f*, or equivalently the complex number *z* = *g* +*is*, the so-called *phasor*, defined by:

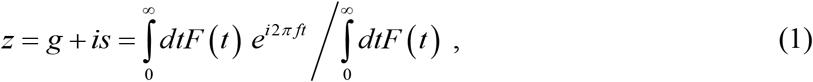

*f*, the *phasor harmonic*, is typically chosen equal to a small integer multiple of the laser repetition frequency. The (*g*, *s*) pair is then represented in a two-dimensional plot, the so-called *phasor plot*, whose properties are summarized in figure 1 and in supplementary note 1. A particularly useful property of the phasor representation is that the phasor *z* of a single-exponential decay is located on the so-called *universal circle*, with short lifetimes at one end (*τ* = 0 → *z* = 1) and long lifetime at the other (*τ* = ∞ → *z* = 0). All other single exponential decays are located in between, as expressed by SI equation (SI-7). Due to the linearity of the Fourier transform, the phasor of a mixture of species is equal to a linear combination of their individual phasors, the weight factor of each species being equal to its emitted intensity (SI equation (SI-18))[19]. The phasor representation of an image, which comprises the phasor of each pixel, is easily computed and represented graphically as a 2-dimensional histogram in the phasor plot. Such a representation allows visual identification of molecular species characterized by distinct fluorescence decays, without any assumption about the underlying decay model function. Following identification of different species of interest in the phasor plot (figure 1), the corresponding sources (*i.e.* image pixels) can be immediately located in the original image. Using color-coding, each identified species can be mapped in a false-color overlay image, allowing rapid identification of a sample’s lifetime heterogeneity. Finally, precise quantification of the respective fractions of two or more components can be performed, provided the location of known species’ phasor location can be determined.

**Figure 1:**
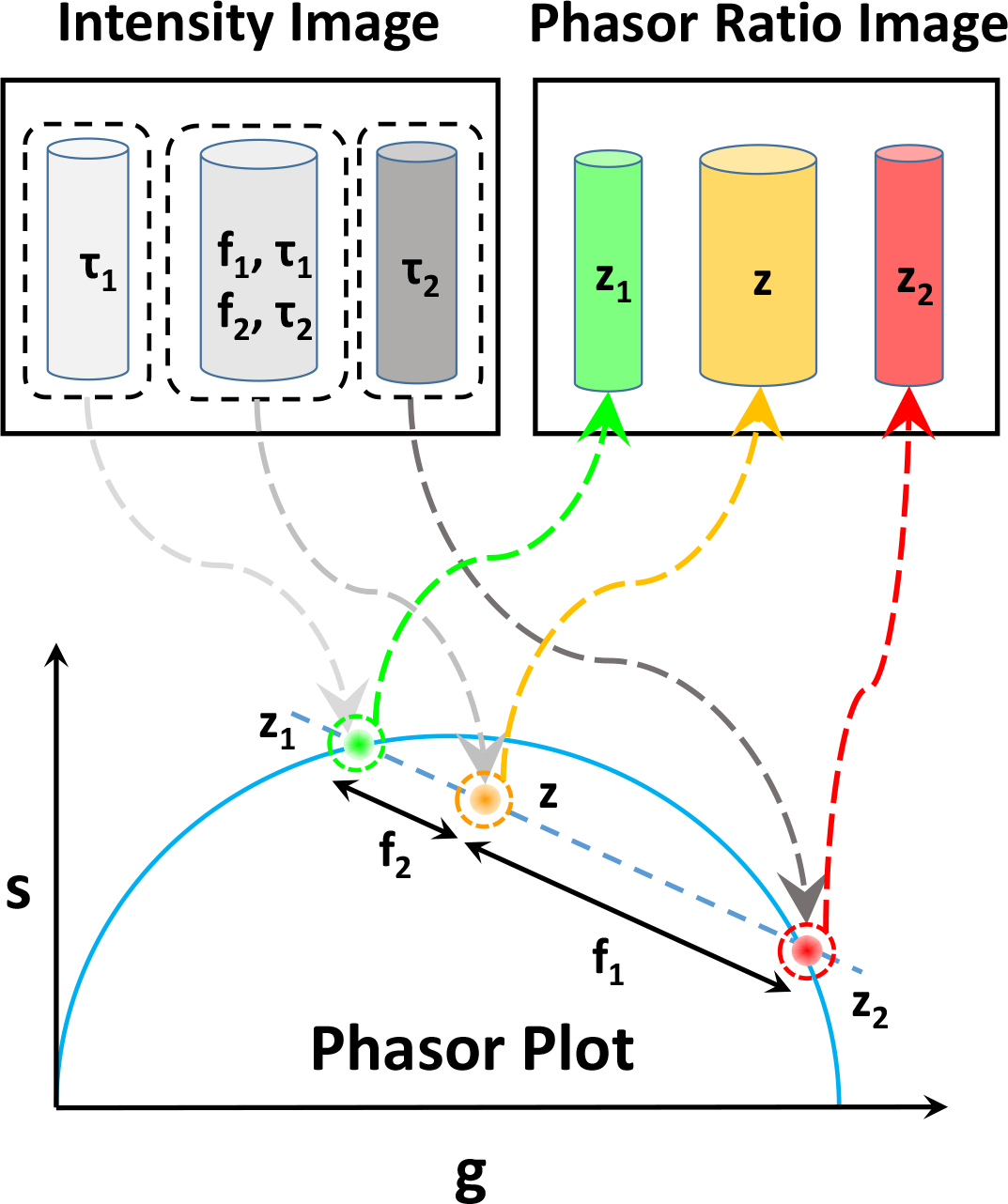
Principle of phasor analysis. Fluorescence lifetime data can be represented with almost no processing as an intensity image (top left panel). The phasor representation of this data (easily calculated for each pixel with equation (SI-1) (supplementary information) can be displayed as a phasor plot (gray arrows and bottom panel), in which pixels characterized by similar fluorescence decays are clustered together around the universal circle (UC, blue), where pure exponential decays are located. This representation then allows the selection of clusters of phasors (e.g. cluster *z*_*1*_: green, *z*_*2*_: red and *z*: orange) and highlighting the corresponding pixels of the image with different colors (colored arrows and top right panel). In the example schematically shown here (top left), samples with lifetime *τ*_*1*_ and *τ*_*2*_, and a mixture of both (fractions *f*_*1*_ and *f*_*2*_), the samples phasors *z*_*1*_ and *z*_*2*_ are located on the UC, while the mixture’s phasor *z* is located on the line connecting them. In this type of cases, it is simple to compute and color-code, almost in real-time, the fraction *f*_*1*_ of sample 1 (also called phasor ratio) in the original image. The resulting “phasor ratio image” is thus a convenient “map” of the fraction of the lifetime of interest within the sample. Quantitative analysis can also be performed, as discussed in the main text and supplementary information.

This method has been successfully applied in a number of *in vitro* studies using fluorophores emitting in the visible range of the spectrum and characterized by fluorescence lifetimes that are long compared to the width of the instrument response function (IRF)[21, 22, 24, 25, 27, 50], including FRET studies[53]. Phasor analysis has also been used to characterize autofluorescence in intravital microscopy-based studies of the brain[32] and kidney[54], as well as in superficial skin layers using two-photon microscopy[26], and for determination of tissue redox ratio[30]. Additionally, the phasor technique has been extended to the spectral domain for characterization of spectrally-dependent lifetime shifts[33] as well as for other imaging applications such as extraction of collagen fibril anisotropy through second harmonic generation[31] and quantitative T2 mapping for magnetic resonance imaging[55].

In this work, we demonstrate that phasor analysis can be successfully used in the challenging situation presented by mixtures of short lifetime NIR dyes, even in the presence of the broad IRF encountered during *in vivo* experiments. We first demonstrate that small NIR lifetime changes (20 ps) can be readily detected and quantified by phasor analysis, even though the intrinsic lifetimes of these dyes are very short. We next build on this result to show that mixtures of short NIR lifetimes can be quantified *in vitro*, using first a pair of spectrally-similar dyes with distinct lifetimes mixed at different ratios. We then extend this demonstration to a single dye exhibiting different lifetimes due to FRET, in two different *in vitro* FRET-based binding assays. Finally, we extend our demonstration to *in vivo* studies by studying the kinetics of transferrin engagement with the transferrin receptor in live mice. We conclude that the quantitative nature, ease of use and interpretation, as well as rapid data acquisition and analysis afforded by the phFLI approach, make it a very powerful new molecular imaging method for *in vitro* as well as *in vivo* real time kinetic studies.

## 2 Materials and Methods

### Buffer mixtures

As an initial test of phFLI for determining fluorescence lifetime, we mixed AF700 (5 μL of a 37.5 μM solution in PBS) with either PBS or ethanol and then added aliquots of each solvent at regular intervals, while measuring the fluorescence of the sample in the meantime (see supplementary table 1 for details of the dilution steps). For the short (PNN) and long (ENN) lifetime control wells in which the lifetime was not modulated, 5μL of AF700 was mixed with 200 μL of PBS, or 200 μL of ethanol respectively. For the other wells, two modulation series consisting of different solvent addition were used. In the first modulation series (starting at data set = 21), 20 μL of the solvent was added every 20 data sets (21 to 81). The second modulation series started at data set = 181, and 20 μL of the next solvent was added every 20 data sets (181 to 301). At the end of each modulation series, the solution in each well was mixed with a pipette tip. Data was collected periodically during and in-between modulation series (total experiment duration: 6,297 s). Data was analyzed by single-exponential decay fitting and phasor analysis as described in supplementary note 1.

### Dye mixtures experiments

To assess the capabilities of phFLI for unmixing two lifetimes, we mixed two NIR dyes, ATTO 740 (A740, 91394-1MG-F, Sigma-Aldrich, MO) and 1,1′,3,3,3′,3′-hexamethylindotricarbocyanine iodide (HITCI, 252034-100MG, Sigma-Aldrich, MO) initially prepared in PBS at various initial concentrations. These dyes were selected for their similar excitation and emission spectra, but distinct lifetime values. For each concentration pair, different volumes of both dyes were mixed to obtain a total volume of 300 μL with volume fractions ranging for 0 to 100% (10% steps, see supplementary table 2). Time-gated imaging was performed with a more sensitive time-gated system than for the other experiments. Data analysis (global lifetime fitting and phasor analysis) was performed as described below. Lifetimes quoted in the text are the mean and standard deviations of the lifetimes fitted in the global analysis of each data set. The lifetimes of the phasor references used in phasor analysis were only slightly larger (A740: 597 ± 34 ps, HITCI: 398 ± 25 ps) by approximately 30-40 ps.

### IgG solutions donor AF700 and acceptor AF750 at various A:D ratios

In order to comprehensively analyze the capabilities of our phasor tools for quantification of NIR lifetimes and FRET, an *in vitro* experiment was designed using pairs of primary and secondary antibodies. We used Alexa Fluor 700 conjugated to murine IgG primary antibody (AF700, MG129, ThermoFisher Scientific, MA) as donor and Alexa Fluor 750 conjugated to goat anti-mouse IgG secondary antibody (AF750, A-21037, ThermoFisher Scientific, MA) as acceptor. The dyes were first diluted to a concentration of 50 μg/mL in phosphate buffered saline (PBS), and then mixed in various acceptor to donor (A:D) ratios (0:1, 1:1, 2:1 and 3:1) in a 96-well microplate to a final volume of 200 μL in each well. The concentration of the donor was kept constant at 40 μg/μL, while that of the acceptor varied with A:D ratio. The well plate was imaged in the reflectance geometry in the FLI system and using the appropriate emission filter sets. The wells were analyzed as individual regions of interest in the bi-exponential fitting analysis to extract the short-lifetime fraction. For phasor analysis, the 0:1 well was used as the donor-only mono-exponential lifetime calibration (*τ*_*D*_ = 1 ns).

### Preparation of fluorescently-labeled transferrin (Tf) conjugates

Human iron-saturated holoTf (T8158, Sigma-Aldrich, MO) was conjugated to Alexa Fluor 700 or Alexa Fluor 750 (A20010 and A20011; ThermoFisher Scientific, MA) according to manufacturer instructions. Briefly, Tf was incubated in 100 mM NaHCO_3_ buffer pH 8.3 with the dyes for 1 hr, followed by passage over a resin desalting column to separate conjugates from free dye. Tf probes were stored in PBS without preservatives. The average degree of labeling was 2 fluorophores per Tf molecule.

### Cell lines

The human breast cancer cell line T47D (HTB-133) was purchased from ATCC (Manassas, VA) and used within one year of purchase. No cells authentication was performed. Cells were cultured in Dulbecco’s modified Eagle’s medium (11965; ThermoFisher Scientific, MA) supplemented with 10% fetal calf serum (30-2020, ATCC), 4 mM L-glutamine (ThermoFisher Scientific, MA), 10 mM HEPES (Sigma) and Penicillin/Streptomycin (50 Units/mL or 50 μg/mL, ThermoFisher Scientific, MA) at 37 °C and 5% CO_2_. Cells were cultured no longer than passage 20 after thawing. Mycoplasma test was performed once in three months using Universal Mycoplasma Detection kit (ATCC).

### Cellular internalization of FRET probes

T47D cancer cells were plated on 60 mm^2^ plastic dishes and cultured until confluent, washed and pre-cleared with serum-free DHB imaging medium [phenol-free Dulbecco’s Modified Eagle Medium (DMEM) (ThermoFisher Scientific, MA), 0.5% bovine serum albumin (Sigma-Aldrich, MO), and 20 mM HEPES (Sigma-Aldrich, MO) pH7.4] at 37 °C for 30 min. Cells were then incubated for 1 hr at 37 °C with the various solutions of Tf-AF700 and Tf-AF750 conjugates diluted in DHB medium at specific Acceptor to Donor (A:D) ratios keeping Donor (Tf-AF700) concentration constant at 40 μg/mL. After incubation, cells were washed with phosphate buffer saline (PBS), fixed with 4% paraformaldehyde, scraped, pelleted, and transferred to a 96-well plate for imaging. The well-plate was then imaged in the FLI system in the transmission geometry. A piece of diffusing white paper was placed beneath the plate on the imaging stage to decrease reflection of stray light, and active illumination was employed to ensure matching of dynamic range between the wells. After imaging, the wells were analyzed separately using the fitting approach to extract the short lifetime fraction. The 0:1 well was used as calibration in the phasor plot, and a circular ROI was used for further analysis.

### In vivo FRET dynamics

In these experiments, we observed the dynamics of FRET by injecting Tf probes labeled with donor and acceptor at different time points. For all experiments, athymic nude female mice (Charles River, MA) were first anesthetized with isoflurane (EZ-SA800 System, E-Z Anesthesia), placed on the imaging stage and fixed to the stage with surgical tape (3M Micropore) to prevent motion. A warm air blower (Bair Hugger 50500, 3M Corporation) was applied to maintain body temperature. The animals were monitored for respiratory rate, pain reflex, and discomfort. The mice were imaged with the time-gated imaging system in the reflectance geometry, with adaptive greyscale illumination to ensure appropriate dynamic range between the regions of interest[56]. In particular, excitation intensity had to be reduced in the urinary bladder due to accumulation of labeled Tf over time. 2 hr after tail-vein injection of 20 μg of Tf-AF700, the mice were imaged for ~15 minutes before 40 μg of Tf-AF750 (A:D ratio = 2:1), and for one of the mice, 100 μg of unlabeled Tf (A:D:U ratio = 2:1:5), were retro-orbitally injected into the other eye. Imaging was continued for another 90 minutes. For the control mouse (0:1), no further probe was injected throughout the imaging session.

### Data acquisition

The time-resolved FLI-FRET imaging system used in this study was described in detail in [57]. Briefly, a Ti:Sapphire pulsed laser (MaiTai HP, SpectraPhysics, CA) is used as an excitation source, delivering 100fs pulses at an 80MHz repetition rate. This laser is tunable from 695-1100nm, which is within the desired NIR range. The power of the laser was controlled by a variable attenuator (CCVA-TL; Newport, CA) to ensure stability. The system is capable of operating in both transmission and reflection modes, and the mode is selected according to the application. For all experiments herein, the samples were excited at the appropriate wavelength and the light is collected by a gated intensified CCD camera (PicoStar HR, LaVision, Germany). The exposure time and multichannel plate (MCP) voltage of the camera are adjusted accordingly in order to optimize the collected signal. For all experiments, the exposure time was in the range 100-800 ms while the laser power was typically below 500 mW. Active illumination[56, 58] was used to further optimize the dynamic range between different regions of interest (ROIs) by changing the spatial patterns projected onto the imaging plane using a digital micro-mirror device (DLi4110; Digital Light Innovations, TX) coupled with the laser excitation. This allows for imaging of multiple regions of interest in one field of view without compromising image quality due to saturation or low signal. For all data sets, an instrument response function (IRF) of the system was collected by imaging the excitation laser light scattered by piece of white paper without emission filter. In practice, we used a deconvolved IRF for *in vitro* data analysis, the “paper” IRF being used for *in vivo* data analysis. The gate width of the camera was set to 300 ps and the gate step to 40 ps. Even though longer gate widths can be used to increase signal without changing the estimation accuracy of FRET parameters[59], a 300 ps gate width was used to allow standard FLI analysis. Two emission filters (FF01-715/LP-25 and FF01-720/13-25; Semrock Inc., NY, for the AF700-based experiments and FF01-820/12-25; Semrock, Inc., NY and ET810/90; Chroma Technology, VT, for the HITCI/A740 mixture experiment) were used to collect the fluorescence signal and reject scattered laser light and, in the case of FRET studies, the acceptor emission.

### Data analysis

Details of data analysis are discussed in supplementary notes 1-4.

### Ethical statement

All animal procedures were conducted with the approval of the Institutional Animal Care and Use Committee at both Albany Medical College and Rensselaer Polytechnic Institute. Animal facilities of both institutions have been accredited by the American Association for Accreditation for Laboratory Animals Care International.

### Software & Data availability

Data and software are available online as described next.

### Software

A Microsoft Windows 64 bit software (AlliGator) developed for phasor analysis and lifetime fitting of time-gated FLI data is provided as a free executable file at https://sites.google.com/a/g.ucla.edu/alligator/installation. All analyses described in the paper were performed with this software. Implementation details are available from the corresponding author upon request. Some of the lifetime fitting results were verified using custom Matlab code from the Intes group, available upon request. Additionally, some of the model calculations presented in supplementary notes 3 & 4 were performed using a separate software (phFRET), available as a Windows executable at https://sites.google.com/a/g.ucla.edu/alligator/phfret-software.

### Data

Datasets and associated acquisition and analysis files, as well as short content description files, are available on Figshare as follows:

- Figure 2 & Supplementary Figure 1: https://doi.org/10.6084/m9.figshare.5561872
- Figure 3 & Supplementary Figure 2: https://doi.org/10.6084/m9.figshare.5776890
- Figure 4: https://doi.org/10.6084/m9.figshare.5786694
- Figure 5: https://doi.org/10.6084/m9.figshare.5788128
- Figures 6 & 7 and Supplementary figures 3-6: https://doi.org/10.6084/m9.figshare.5791476

**Figure 2:**
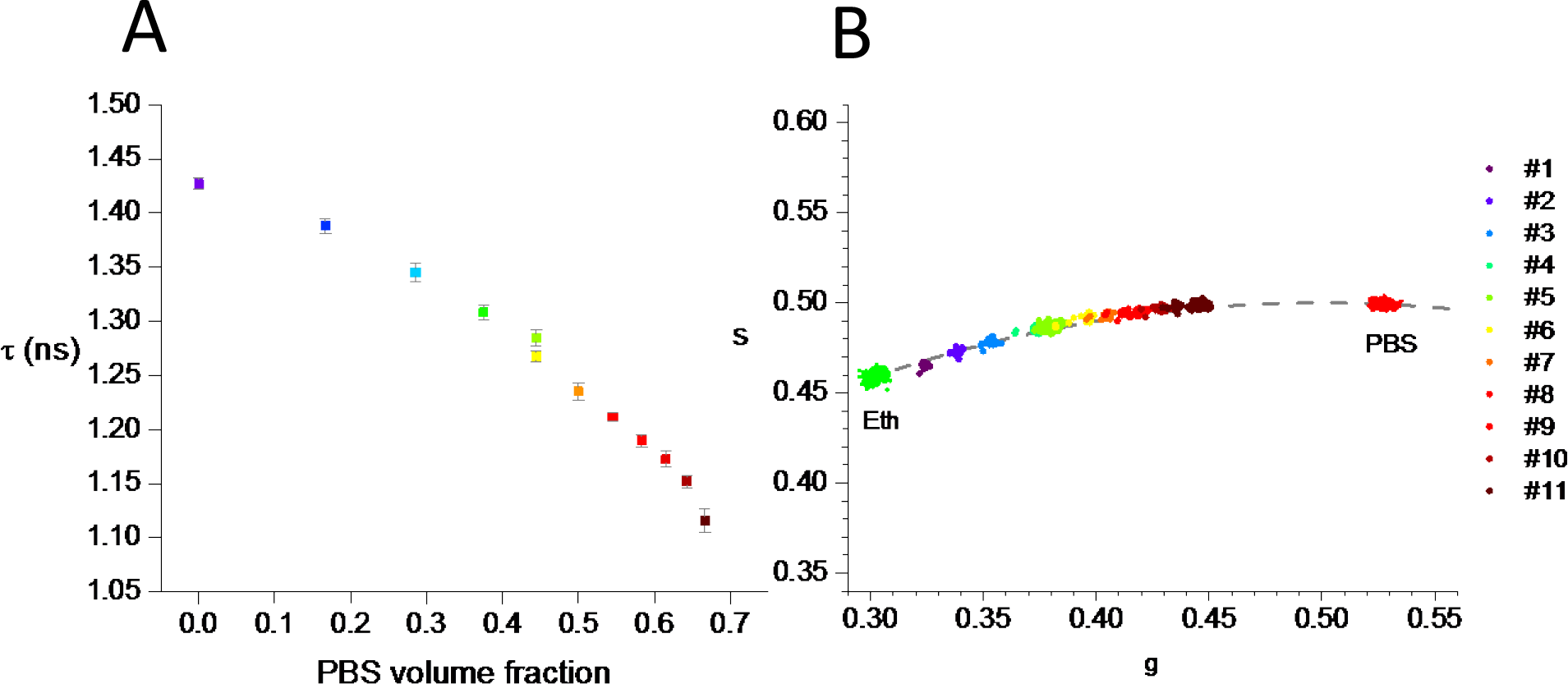
Phase lifetime (A) and phasor (B) of AF700 in EtOH:PBS as a function of PBS volume fraction. A: Each point corresponds to the average of 20 measurements (*). The error bar shows the standard deviation for each mixture. The measurable changes in lifetimes from one condition to the next range from 17 to 43 ps, with a typical standard deviation within each measurement of 10 ps. The instrument response function’s FWHM was 177 ± 7 ps. (*) Points 5+6 and 11 correspond to the average of 100 measurements. B: Detail of the phasor graph showing the corresponding phasors for each individual measurement (noted #n) as well as for the samples in pure ethanol (left, green) and pure PBS (right, red), used as reference for calibration. Clusters for each dilution are readily separated. Some small temporal evolution of the sample is clearly visible during each series of measurements. Additional details in online Methods, supplementary figure 1 and supplementary movie 1.

**Figure 3:**
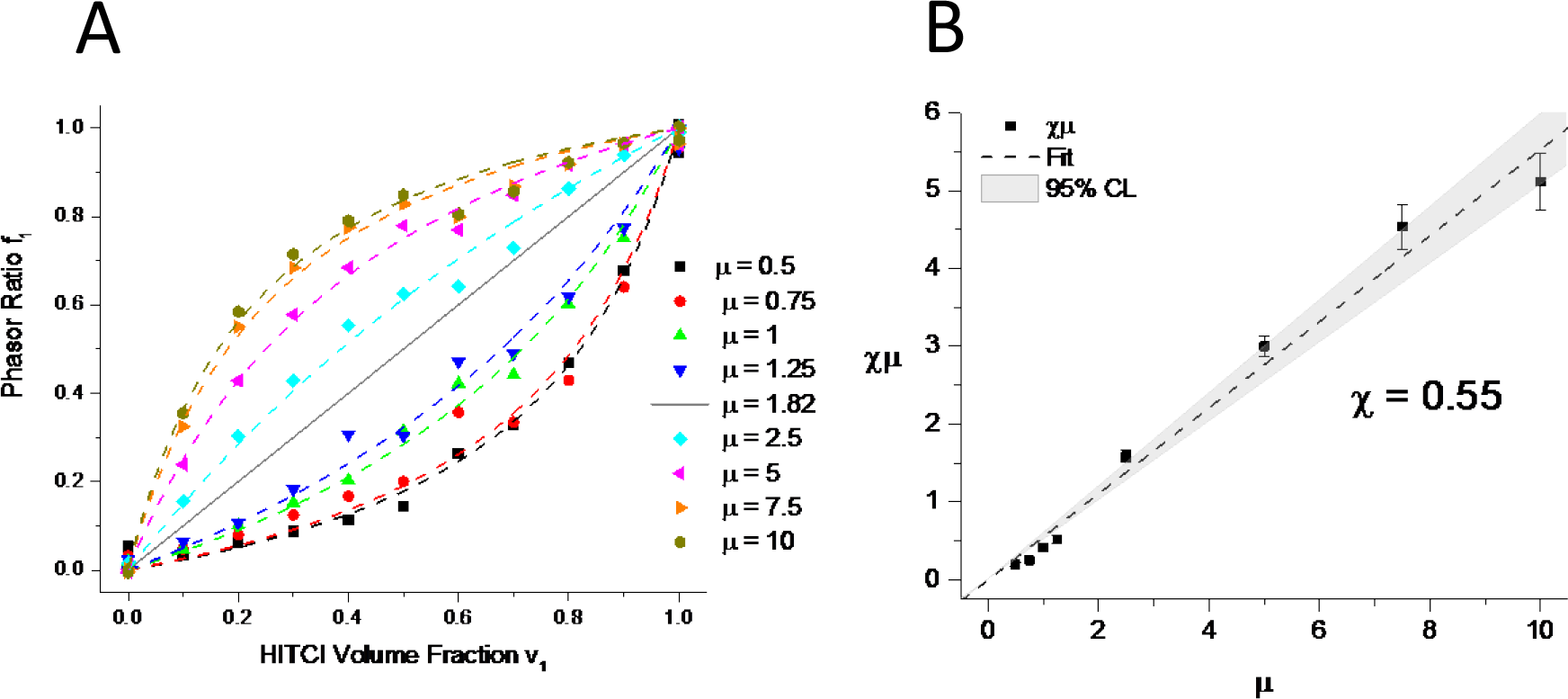
Analysis of series of mixtures of dyes with different lifetimes. A: Phasor ratio of different series of mixtures of ATTO 740 and HITCI. Each series is characterized by a variable volume fraction *v*_*1*_ of HITCI, *v*_*1*_ = 0, 0.1,…, 0.9, 1 and initial concentration ratio *μ* = [HITCI]_0_/[A740]_0_ given in the legend. Each data series is fitted with the theoretical expression for *f_1_(v_1_)* discussed in the text, yielding a parameter value *χμ* studied in B. B: Relation between *χμ* (obtained from the analysis described in A) and *μ*. A linear dependence is verified, with *χ* = 0.55.

**Figure 4:**
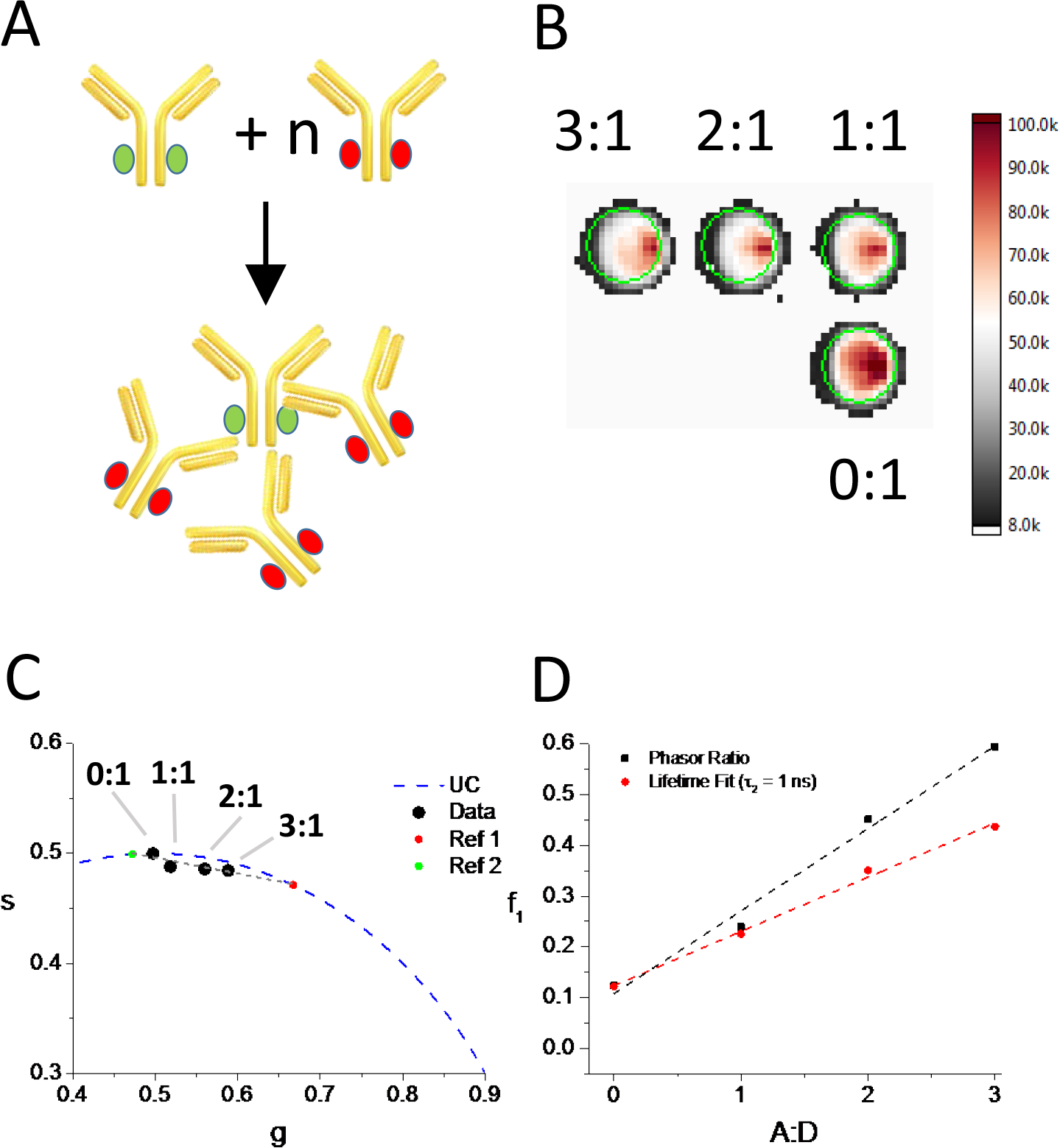
*in vitro* analysis of IgG pair interaction. A: Principle of the experiment. N-fold amounts of acceptor-labeled secondary IgG (red) are added to a fixed amount of donor-labeled primary IgG (green), to form secondary/primary IgG complexes. B: Resulting samples are analyzed with time-gated imaging (shown are the total intensity images, summing all time-gate images). C: Phasor analysis of each sample, once calibrated with the known lifetime of the pure donor sample (0:1), shows alignment of the different phasors. The two limiting values correspond to the pure donor sample (0:1, green), and a hypothetical, fully quenched sample (red) with lifetime *τ*_*DA*_ = 703 ps). D: Representation of the phasor ratio *f*_*1*_ (black squares) for these samples, demonstrating a linear dependence on the A:D ratio, in reasonable agreement with the results obtained by lifetime fitting (red circles).

**Figure 5:**
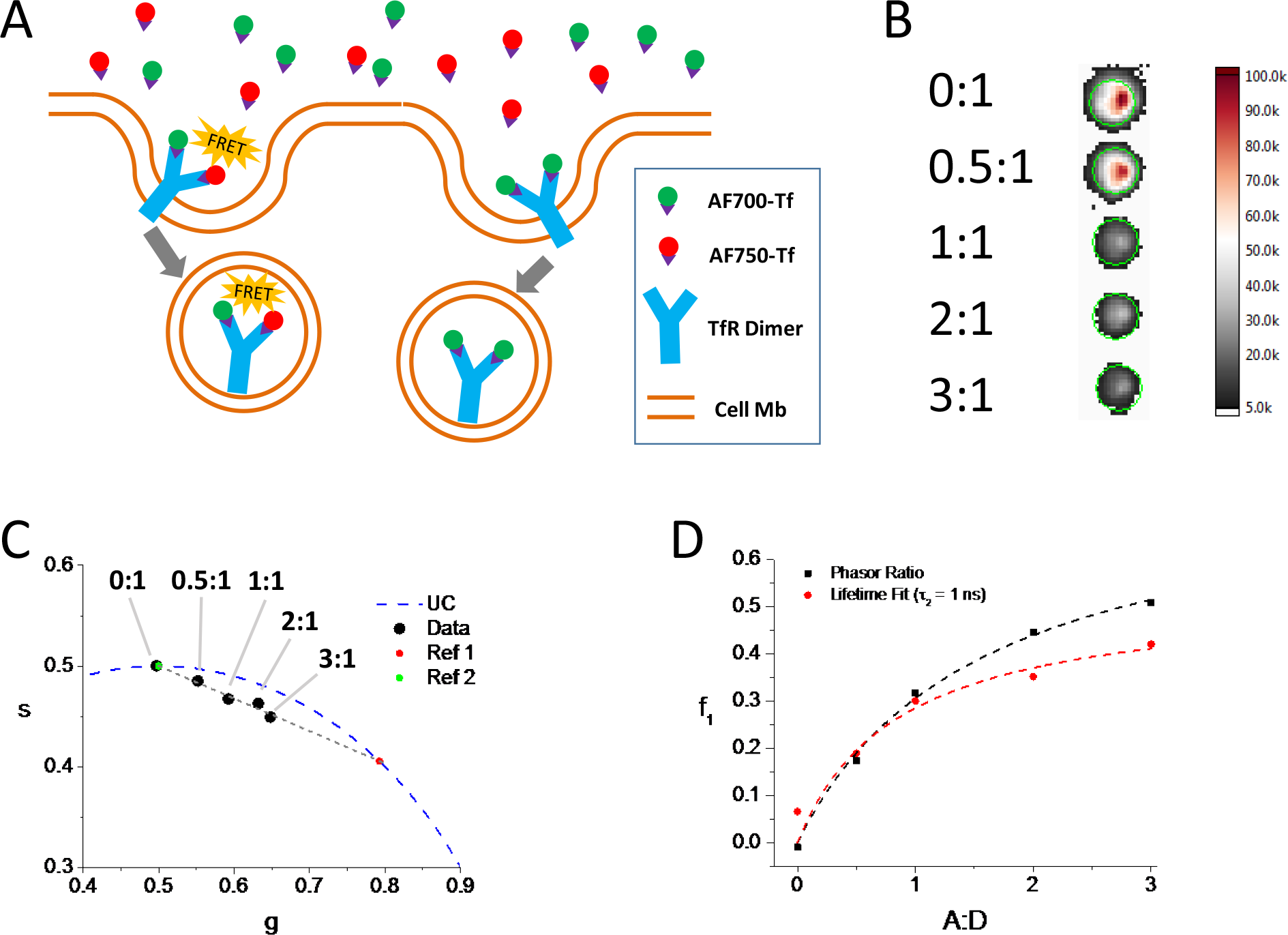
*in vitro* analysis of transferrin receptor (TfR) engagement by transferrin (Tf). A: Principle of the experiment. Different amounts of acceptor-labeled (red) or donor-labeled Tf (green) were added to T47D cells expressing TfR on their surface, leading to Tf binding and TfR internalization by endocytosis. B: Total intensity images of fixed T47D cell cultures incubated with varying ratios (A:D) of acceptor-to donor-labeled Tf. C: Phasor analysis of each sample, once calibrated with the known lifetime of the pure donor sample (0:1), shows alignment of the different phasors. The two limiting values correspond to the pure donor sample (0:1, green), and a maximally quenched donor (red) with lifetime *τ*_*DA*_ = 509 ps). D: Representation of the phasor ratio *f*_*1*_ showing increase and saturation as a function of A:D ratio (black square). The corresponding quantity obtained by lifetime fitting analysis is shown for comparison (red circle). Both agree qualitatively but differ due to the different lifetimes extracted by global fitting of the dataset (*τ*_*DA*_ = 391 ps when using *τ*_*D*_ = 1 ns).

**Figure 6:**
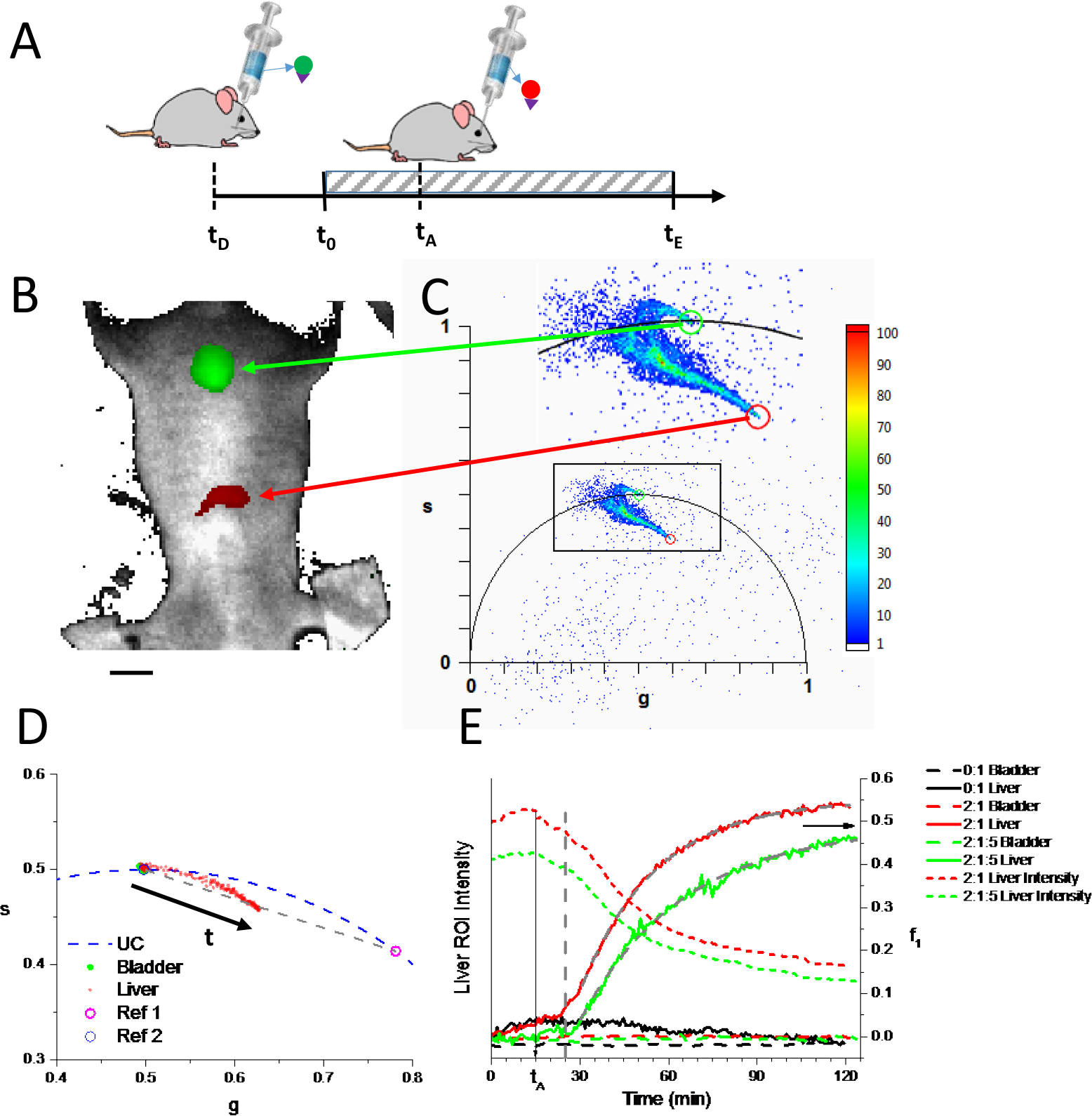
*in vivo* kinetics of transferrin engagement with Tf-receptor. A: principle of the experiment. Mice were first injected with Tf-AF700 (time tD), and imaging started after several minutes (time t0). 15 min after the beginning of the imaging session, Tf-AF750 was injected (time tA) and imaging continued for another 1.45 hr. B: White light image of an anesthetized mouse (2:1:5) about half-way through the imaging session. The bladder (top, green) and liver (bottom, red) are highlighted by selection of two localized regions in the phasor plot shown in C. C: Phasor plot (and detail of the boxed area), with two rectangle ROIs selecting particular regions of the plot. D: Phasor plot showing the loci of the bladder ROI’s average phasor (green dots) and liver ROI’s average phasor (red dots) over the duration of the whole experiment. The references used for phasor ratio calibration are indicated as Ref 1 (0.53 ns) and Ref 2 (1 ns). E: Liver ROI intensity (left) and phasor ratio f_1_ (right) for (1) a control mouse without Tf-AF750 injection (0:1: black), (2) a mouse injected with 2:1 Tf-AF750 injection at t_A_ = 15 min (2:1: red) and (3) a mouse injected with 2:1 Tf-AF750 and 5:1 unlabeled Tf at t_A_ (2:1:5: green). Short dashed curves represent the liver ROI intensity, dashed curves correspond to the bladder phasor ratio, while plain curves of the same color correspond to the liver phasor ratio. Gray dashed curves are exponential fits of the experimental curves (for t > 30 min) with τ_2:1_ = 1,528 ± 15 s and τ_2:1:5_ = 2,109 ± 55 s. The vertical arrow indicates t_A_. The vertical dashed line is a guide for the eye indicating the onset of phasor ratio increase. See supplementary movie 2 for an animation of panel B &C. A detailed discussion of the influence of the phasor dispersion observed in C on the analysis, as well as details on how the liver phasor trajectory shown in D is calibrated, can be found in supplementary note 2, and is illustrated in supplementary figures 3 & 4.

**Figure 7:**
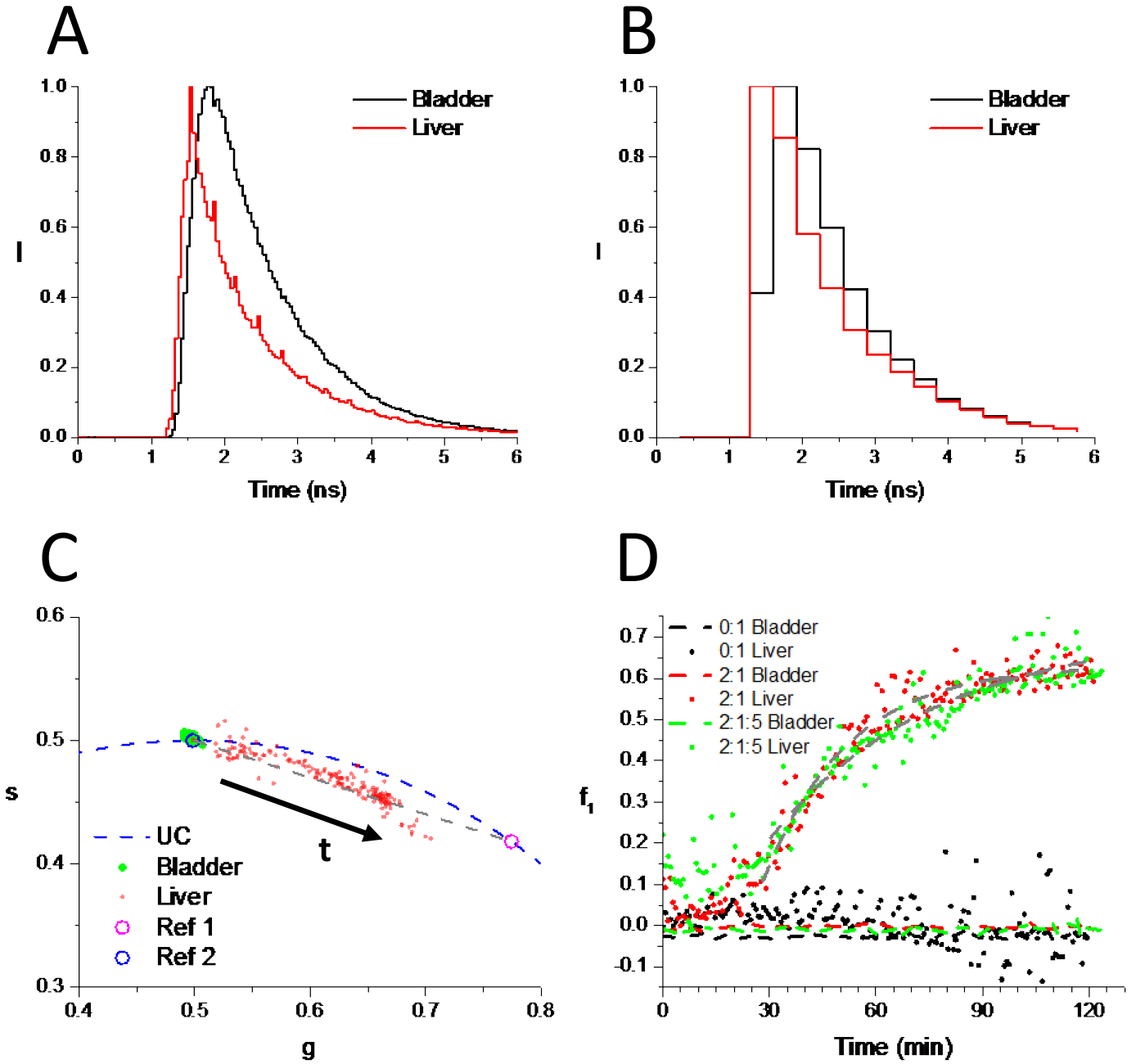
*in vivo* kinetics of transferrin engagement with Tf-receptor using a reduced number of gates. A: Example of full decay (168 gates) obtained in the bladder (black) and liver (red) of mouse 2:1:5. The spikes in the liver decay are due to the mouse’s 0.7 Hz breathing pattern and reflect the sensitivity of the signal to source depth. B: Same decays as in A, represented with a subset of 20 contiguous gates. These coarse resolution decays were used for the remainder of the analysis. C: Detail of the corresponding phasor plots. Compare with the phasor plot obtained using all gates in Fig. 6D. Ref 1 and 2 where defined using the reprocessed data. D: Bladder and liver phasor ratio (f_1_) plots for 3 mice. Dashed curves correspond to the bladder’s phasor ratio, while dotted curves of the same color correspond to the liver phasor ratio. Gray dashed curves are exponential fits of the experimental curves (for t > t_A_) with τ_2:1_ = 1,418 ± 93 s and τ_2:1:5_ = 2,769 ± 356 s. Notice that the asymptotic values are different than those obtained with the full set of gates (Fig. 6E), due to the different definitions of the short lifetime intersection (reference 1). Using the same references in both analyses would have resulted in similar asymptotic values. Additional discussion of the difference between phasor analysis with a reduced number of gates and the full set of gates can be found in supplementary note 2 and supplementary figure 5.

## 3 Results and Discussion

### 3.1 ACCURATE MEASUREMENT OF SHORT NIR LIFETIMES

We first characterized the ability of NIR phFLI to precisely quantify the short fluorescence lifetimes which are typical of most NIR dyes, by taking advantage of the exquisite sensitivity of lifetime on specific environmental perturbations. Since we used it in most of our experiments, we characterized Alexa Fluor 700 (AF700), which has a reported lifetime of 1 ns in water. Like many dyes, such as dyes from the oxazine family, AF700 exhibits a larger lifetime in organic solvents (for instance ethanol, EtOH) than in aqueous buffers (for instance phosphate buffered saline, PBS)[60, 61]. The lifetime of AF700 can thus be modulated in a predictable manner by changing the ratio of EtOH to PBS in the solution. Figure 2A-B and supplementary movie 1 show the changes in the phase lifetime of AF700 (supplementary information (SI) equation (SI-7)) and phasor as different amounts of PBS were added to a 100 μL EtOH solution of AF700. Lifetime steps of 20 ps or less are clearly identifiable. Similar results were obtained using different starting solutions and dilution sequences (supplementary figure 1A-B), consistent with quenching of AF700’s fluorescence by PBS (supplementary figure 1C). Importantly, the extracted phase life-times (obtained by phasor analysis) were comparable to the lifetimes obtained by a standard single-exponential fitting approach (supplementary figure 1D), although they were performed completely independently from one another.

### 3.2 ANALYSIS OF MIXTURES OF NIR SPECIES CHARACTERIZED BY DIFFERENT LIFETIMES

Having established the ability of phFLI to measure short fluorescence lifetimes of individual NIR dyes, we studied *mixtures* of NIR dyes characterized by *similar* emission spectra but *different* lifetimes and asked how reliably the *fraction* of each species can be measured by phFLI. The ability to quantify the amount of fluorophores differing only by their lifetime would allow using lifetime as a contrast mechanism, and open up the possibility to use it to analyze molecular interactions by FRET.

In traditional lifetime fitting approaches, assuming that each component of a mixture is characterized by a distinct, single exponential decay, a simple definition of the fraction of each component is given by the relative amplitude *α*_*i*_ of each exponential component of the mixture’s decay (SI equation (SI-24)). Alternatively, the phasor *z* of a mixture of different species, which is simply a weighted sum of each individual species’ phasor *z*_*i*_ (SI equation (SI-18)):

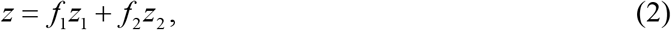

leads to a natural definition of each species’ fraction as the phasor weight *f*_*i*_ of this species. As explained in Figure 1 and in supplementary note 1, if the phasors of each isolated species are known (for instance from separate measurements of each individual dye), *f*_*i*_ is given by the relative distance of the sample’s phasor to that species’ phasor (the so-called *phasor ratio*, SI equation (SI-20)-(SI-21)):

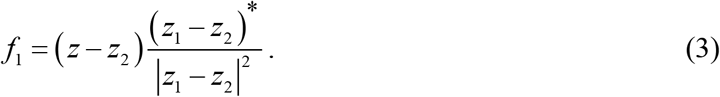

*α*_*i*_ and *f*_*i*_ can be used interchangeably if the two lifetimes of the reference species are known (SI equation (SI-25)). These quantities can be connected to independently measurable observables such as the species’ *volume fraction v* (SI equation (SI-35)), *number fraction n* (SI equation (SI-37) or *concentration ratio M* (SI equation (SI-33)), by simple expressions involving a single parameter *χ* characterizing the dyes’ photophysical properties and instrument sensitivity (SI equation (SI-30)) and the initial concentration ratio *μ* of the two dyes (SI equation (SI-34)):

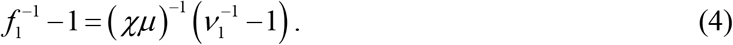

In particular, *μ* can be chosen such that *f*(*v*) becomes linear.

Figure 3 shows the results of phasor analysis of different mixture series of various initial concentrations of ATTO 740 (A740) and 1,1′,3,3,3′,3′-Hexamethylindotricarbocyanine iodide (HITCI), two NIR dyes which can both be excited at 740 nm and emit around 770 nm, and are characterized by different lifetimes in PBS buffer (A740: 567 ± 11 ps, HITCI: 356 ± 4 ps). Each series corresponds to variable volume fractions *v*_*1*_ of HITCI (species 1, initial concentration [HITCI]0) and A740 (species 2, initial concentration [A740]_0_), and is fully characterized by the ratio *μ* of their initial concentrations. As predicted by SI equation (SI-35), the phasor ratios of each series depend only on *μ* and the volume fractions *v*_*1*_, the dependence becoming linear for a single value *μ* = χ-1. Importantly, the lifetime fitting analysis performed independently provides identical results (supplementary Figure 1).

In other words, knowing *χ* and the lifetimes of the two dyes, it is possible to recover the volume fraction in any mixture based on the measurement of the phasor ratio *f*. These results demonstrate the ability of phFLI analysis to provide quantitative information on the composition of mixtures of species characterized by similar emission spectra (and therefore undistinguishable spectroscopically) but different lifetimes.

### 3.3 ANALYSIS OF *IN VITRO* NIR FRET

The ability to distinguish identical dyes differing only by their lifetime is, in particular, useful for the analysis of binding equilibria by FRET. In this application, the association of two molecules, one of which is labeled with a donor dye and the other with an acceptor dye, is detected by energy transfer of the donor to the acceptor. In the absence of binding, no FRET occurs and the measured donor lifetime is simply the unquenched donor lifetime *τ*_*D0*_. In the case of binding, FRET between donor and acceptor results in a decreased lifetime *τ*_*DA*_. If the bound donor-acceptor configuration is unique, the FRET efficiency (hence the FRETing donor lifetime *τ*_*DA*_) is a constant and *τ*_*DA*_ can be considered as the second reference lifetime of the mixture of bound and unbound donor molecules. Measurement of the fractions of the fluorescence decay with lifetime *τ*_*D0*_ or *τ*_*DA*_ is then equivalent to measuring the respective fraction of unbound and bound donor respectively, and thus allows full characterization of the binding equilibrium. This can be challenging to do using multi-exponential fits, in particular when short lifetimes are involved, but as discussed in the next two examples, this is straightforward using phFLI analysis.

#### 3.3.1 Quantification of FRET between IgG antibodies

In the first example, schematically depicted in figure 4A, known amounts of donor-labeled mouse primary antibody (AF700-IgG) were mixed with increasing amounts of acceptor-labeled goat anti-mouse secondary antibody (AF750-IgG). When increasing the acceptor-labeled to donor-labeled IgG ratio (A:D), the average number of AF750-IgG molecules bound to each AF700-IgG molecule increases, resulting in more FRET interactions and thus a larger fraction of donor molecules involved in FRET. As discussed above, these changes are reflected in the donor fluorescence decay, and have previously been measured by time-gated fluorescence imaging and lifetime fitting analysis[57]. Here, time-gated imaging data was analyzed by phFLI and compared to the corresponding fluorescence decay fitting results.

Figure 4C represents the series of phasor locations for these different A:D ratios. They are clearly aligned along a segment connecting the donor-only phasor located on the universal semicircle (green dot), and another intersection with the semicircle (red dot), identified with the hypothetical maximally quenched donor. Figure 4D shows the observed linear dependence of the phasor ratio as a function of the A:D ratio (black squares), as predicted by the simple model discussed in supplementary note 3, despite the multiplicity of lifetimes involved in this system. This result agrees qualitatively well with the corresponding short lifetime fraction extracted by standard bi-exponential lifetime fitting analysis (red circles), which is not expected to perform well in the presence of potentially more than two lifetimes, showing that phFLI can quantitatively report on the equilibrium association between FRET binding partners *in vitro*.

#### 3.3.2 Quantification of FRET receptor-ligand binding in cells

A particularly important example of molecular interactions that can be studied by FRET methods is that of ligand-receptor binding in cells. Many such interactions are implicated in signal transduction pathways, and represent a major target of drug development [62]. Although labeled antibodies (such as used in the experiments discussed above) can be used for cell studies of ligand-receptor binding[63], a more effective approach available when receptor dimers exist, consists in using labeled ligands. By labeling a fraction of the externally delivered ligands with a donor molecule (D) and another fraction with an acceptor molecule (A), it is then possible to quantify the fraction of doubly-labeled (D + A) receptors by FRET. As in the previous example using labeled antibodies, measurements can be done by time-gated donor lifetime measurements and analyzed by phFLI (figure 5 and supplementary note 4 for theoretical details).

In a large range of situations, the predicted doubly-labeled equilibrium fraction depends simply on the initial A:D ratio. phFLI analysis of donor fluorescence then enables plotting the expected phasor of the mixture as a function of the initial A:D ratio and, with appropriate reference values, the FRET fraction. Two distinct regimes can be identified: (i) one in which the FRET fraction increases to asymptotically reach a maximum value, and (ii) another where the FRET fraction varies linearly with the A:D ratio. Both regimes are of practical relevance: the first regime is that observed in *in vitro* cell measurements discussed in this section, while the second regime is relevant for the *in vivo* experiments described in the next section.

Here, we used holo-transferrin (Tf), an iron-binding blood plasma glycoprotein commonly used as drug carrier[64], and the transferrin receptor (TfR1), a receptor typically overexpressed in cancer cells and the liver. T47D breast cancer cells were incubated for 1 hr with donor- and acceptor-labeled holo-Tf (Tf-AF700 and Tf-AF750) at different A:D ratios. After fixation, the cells were imaged with the time-gated FLI system and analyzed using both phFLI and standard decay fitting approaches. Figure 5C shows the phasor locations for different A:D ratios which, as in the previous experiment, appear aligned on a segment connecting the donor-only sample and an hypothetical maximally-quenched sample. Converting these locations into phasor ratios (or equivalently, FRET population fractions), the curve shown in figure 5D (black squares) is obtained, which confirms the trend obtained by standard decay fitting (red circles), and corresponds to the expected asymptotic saturation predicted by the simple model described in supplementary note 4 (dashed curves).

### 3.4 QUANTIFICATION OF*IN VIVO* FRET DYNAMICS

Having validated the use of phasor analysis with NIR FRET *in vitro*, we examined whether a similar type of analysis would be possible *in vivo*. While *in vivo* NIR fluorescence lifetime imaging is essentially devoid of background autofluorescence, it is complicated by light scattering and absorption by tissues, which broaden the temporal response and attenuate the signal intensity[65, 66]. Since these effects may be depth-dependent, animal respiration, which periodically affects the depth of some organs such as the liver, could potentially complicate the analysis. Finally, a fraction of injected fluorescent molecule is initially non-specifically distributed throughout the body, potentially obfuscating the analysis.

We have previously demonstrated that *in vivo* FLI FRET imaging of intravenously injected donor- and acceptor-labeled holo-Tf in anesthetized mice bearing tumor xenografts allows monitoring TfR engagement by Tf[40]. In these studies, we measured detectable accumulation of doubly-labeled TfR in tumors at 1 hr post-injection based on double-exponential fitting of observed decays. We later used the same animal model system to demonstrate the advantages of adaptive wide field illumination (AWFI) to quantify receptor engagement at 6 and 24 hr post-injection in organs with high endogenous levels of TfR expression such as the liver and kidneys[58]. Here, we focused our study on the early time scale (0-2 hr) post intravenous injection, and performed *continuous* time-gated fluorescence imaging of the injected Tf-AF700, in order to investigate the initial kinetics of non-specific accumulation of Tf in the liver, as well as its elimination in the bladder. Donor-(AF700) and acceptor-(AF750) labeled holo-Tf were injected retro-orbitally in anesthetized mice at distinct time points separated by approximately 15 minutes (figure 6A). Mice injected with an A:D ratio of 0:1 (no acceptor-labeled Tf) were used as control. An A:D ratio of 2:1, determined to be sufficient to observe TfR engagement in our previous studies[40, 58], was used for the other experiments (figure 6). Donor FLI acquisition was performed every ~45 seconds over ~2 hours and data was analyzed by phFLI analysis and standard decay fitting analysis as described in supplementary methods. Figure 6B shows the fluorescence intensity image for one of the 2:1 mice in the middle of the measurement and figure 6C the corresponding phasor plot. It is immediately apparent that the distribution of phasors in a live mouse is more complex than that observed in the previously described *in vitro* experiments. While an exhaustive analysis of the various contribution of local environment, source depth and scattering by tissue is beyond the scope of this study, a qualitative discussion of their magnitude and impact on the following result is presented in supplementary note 2 and supplementary figures 3 & 4. Despite this complexity, it is however straight-forward to find the original locations of specific phasors: phasors located toward the top of the universal circle (green rectangle in figure 6C) originate from the bladder region (green overlay in figure 6B), while phasors located toward the short lifetime region (red rectangle in figure 6C) originate from the liver (red overlay in figure 6B), where overexpression of TfR allows formation of donor-acceptor-labeled Tf pairs resulting in FRET (shorter lifetimes). The remainder of phasor values correspond to other locations in the animal in which circulating Tf-AF700 is found at diverse depth and in various local environments.

As shown in supplementary movie 2, color coding of the different phasor plot areas (long lifetime: green, short lifetime: red) allows qualitatively visualizing the progressive increase of the fraction of Tf-AF700 undergoing FRET with its AF750-labeled counterpart, without any fitting or sophisticated analysis. In addition to this qualitative use of the phasor plot, a quantitative analysis of the FRET fraction present in different regions of the mouse can be performed as in the *in vitro* examples discussed previously. Figure 6D shows the location of the averaged phasors of both bladder (green dots) and liver (red dots) at each time point during the measurement. While the calibrated bladder phasor stays approximately in the same location, the liver phasor slowly migrates towards a region of the phasor plot corresponding to shorter lifetimes. Natural reference points to study this evolution are the calibration phasor (single exponential decay, 1 ns, blue dot) and the extrapolation of the liver phasor “trajectory” to the universal circle (magenta dot). Computing the corresponding phasor ratios (short lifetime fraction, SI equation (SI-20)) for these two phasor populations (bladder and liver) result in curves shown in figure 6E. The mouse injected with donor-labeled Tf only (0:1) exhibits a fairly constant phasor ratio consistent with no significant fraction of Tf-AF700 bound together with Tf-AF750 to TfR. Mice injected successively with donor- and acceptor-labeled Tf (2:1 & 2:1:5), show distinct kinetics, characterized by an exponential increase of the short lifetime phasor component shortly (but not immediately) after injection.

These trends are supported by standard decay fitting analysis (supplementary Figure 6), but are here obtained without the complexity and processing time associated with that latter approach. Interestingly, the recorded donor-intensity in the liver decreases immediately after injection (short dashed curves in figure 6E), while lifetime analysis detects a delayed onset of donor quenching by Tf-AF750.

### 3.5 CHECKING THE SPEED LIMIT OF PHFLI

Having demonstrated the ability to detect subtle changes in molecular targeting in live mice by phasor analysis, we asked whether we could take advantage of the negligible computational cost of phFLI and its robustness with respect to decay temporal resolution[52], to speed up fluorescence lifetime analysis and enable real-time phFLI. To reduce the effective acquisition time, we reprocessed the data presented in Figure 6 using only 1/8 of the time-gate images, keeping only non overlapping gates covering the whole decay (figure 7A). The low resolution fluorescence decays obtained with this reduced number of gates (19 gates) is clearly insufficient for a meaningful analysis by bi-exponential fitting, but sufficient to compute individual pixel phasors and phasor ratios of regions of interest, as illustrated in figure 7B & 7C and supplementary figure 5. These results are qualitatively comparable to those obtained using all 150 gates (figure 6D) & 6E, demonstrating that an acquisition time 8-fold shorter (~4 s per time point) could have been used without significantly affecting phFLI analysis. A further 6-fold speed-up could in principle be obtained by reducing the integration time per gate (190 ms for the data shown in figure 6 & 7), down to the minimum frame duration of the camera (30 ms), as the signal-to-noise ratio of these measurements was very high (single pixel: 47, whole ROI > 400). A 19-gate image could then be acquired in less than 1 s, providing data which could be processed by phFLI and represented in real-time.

## 4 Conclusion

We have established that phasor analysis of time-gated lifetime images can measure minute changes in the lifetime of NIR dyes characterized by short intrinsic lifetimes *in vitro* (~20 ps over lifetimes < 1 ns). We have also verified that phasor analysis is able to quantify the composition of a mixture of fluorophores characterized by single-exponential decays *in vitro*, even when the dyes fluorescence lifetimes are comparable to the width of the instrument function (*τ*_*HITCI*_ = 356 ps, FWHMIRF ~ 250 ps). We then exploited these properties of phasor analysis to quantify molecular interactions by FRET in two *in vitro* experiments, involving respectively fluorescently labeled complementary antibodies in solution, and fluorescently labeled ligands of membrane receptors in cell cultures. The results obtained by phasor analysis were comparable to those obtained by classical fluorescence decay fitting analysis. Finally, we extended these studies to the much more challenging situation of *in vivo* imaging, where fluorescence intensity decays are affected by scattering and absorption in a complex and spatially variable manner. We showed that even in this complex system, phasor analysis can be performed fairly straightforwardly and provides semi-quantitative results in good agreement with those obtained by fluorescence lifetime fitting. Importantly, phasor results were obtained almost instantaneously and robustly, while fluorescence lifetime fitting analysis was both computationally intensive and sensitive to initial conditions.

The demonstration that phasor analysis can be extended to the NIR domain where lifetimes are short and comparable to the IRF width, and in particular to *in vivo* imaging, has many important implications. In addition to a decrease in processing time by several orders of magnitude compared to global multiple lifetime fitting approaches needed for quantitative analysis of lifetime mixtures, the robustness of phasor analysis to coarse-graining will allow reducing the acquisition time and/or excitation intensity accordingly, minimizing laser power requirement and animal or sample exposure, as well as improving the temporal resolution of kinetic measurements. While the system used in this study is limited to frame rates slightly lower than 1 Hz due to camera sensitivity and readout rate, newer models of time-gated cameras using sCMOS sensors and more sensitive photocathodes should be capable of much faster frame rates, only limited by the signal to noise ratio of the recorded time-gate images.

In addition to this speed advantage, phasor analysis of *in vivo* NIR time-gated data also provides a practical approach to implementing multiplexed imaging using lifetime as a contrast mechanism. NIR fluorescent dyes have broad emission spectra that are difficult to spectrally un-mix, due to the complex interplay between wavelength-dependent absorption and scattering in tissues. While fluorescence intensity decays are affected by these tissue characteristics, the measurements presented here demonstrate that these effects can be effectively compensated with prior information on the expected location of individual species in the phasor plot. In particular, the visualization of Tf receptor engagement in the liver illustrated in supplementary movie 2, and performed with no other aid but graphical selection of regions of interests in the phasor plot, shows that two fluorescent species characterized by sufficiently different lifetimes (e.g. 0.5 ns and 1 ns) can be readily separated and quantified. Importantly, such identification does not require fluorescent species to be characterized by single-exponential decays. The combination of imaging speed and life-time contrast capability could be advantageously used for image-guided surgery using NIR markers differing mainly by their lifetimes (e.g. ICG and IRDye-800CW).

The quantitative nature of phasor analysis also suggests that it could be used as a replacement to or as a first estimator for the time- and memory-intensive multi-exponential decay fitting approach used in multiplexed fluorescence lifetime tomography imaging[67, 68]. Finally, while our experiments used a time-gated detection based on intensified camera, the same analysis could be performed with data acquired by time-gated or time-resolved single-photon avalanche diode arrays[69] or time-resolved wide-field photon-counting imagers[23].

## ACKNOWLEDGEMENTS

This work was supported by NIH grant R01 EB19443 (XI & MB). XM was supported in part by DOE grant DE-FC02-02ER63421. The support of Shimon Weiss, in whose lab part of this work was performed, is gratefully acknowledged.

SJC & NS contributed equally to this work. SJC, NS and AR performed experiments. SJC and XM performed phasor analysis. SJC and NS performed lifetime fitting analysis. MB, XI, XM designed the experiments. XM worked on the theory and developed the software. SJC and XM wrote the manuscript draft. All authors contributed to the manuscript. The authors declare no competing interests.

## REFERENCES

[1] M. L. James, S. S. Gambhir, Physiol. Rev. 2012, 92, 897.

[2] C. Kim, C. Favazza, L. H. V. Wang, Chem. Rev. 2010, 110, 2756.

[3] J. V. Frangioni, Curr. Opin. Chem. Biol. 2003, 7, 626.

[4] S. A. Hilderbrand, R. Weissleder, Curr. Opin. Chem. Biol. 2010, 14, 71.

[5] J. K. Willmann, N. van Bruggen, L. M. Dinkelborg, S. S. Gambhir, Nat. Rev. Drug. Discov. 2008, 7, 591.

[6] D. K. Osmonov, D. Heimann, I. Janssen, A. Aksenov, A. Kalz, K. P. Juenemann, Springerplus 2014, 3, 340.

[7] M. S. Ozturk, C. W. Chen, R. Ji, L. L. Zhao, B. N. B. Nguyen, J. P. Fisher, Y. Chen, X. Intes, Ann. Biomed. Eng. 2016, 44, 667.

[8] M. S. Ozturk, D. Rohrbach, U. Sunar, X. Intes, Acad. Radiol. 2014, 21, 271.

[9] F. Danhier, O. Feron, V. Preat, J. Control. Release 2010, 148, 135.

[10] J. Fang, H. Nakamura, H. Maeda, Adv. Drug Deliver. Rev. 2011, 63, 136.

[11] K. Suhling, P. M. W. French, D. Phillips, Photoch. Photobio. Sci. 2005, 4, 13.

[12] M. Y. Berezin, S. Achilefu, Chem. Rev. 2010, 110, 2641.

[13] E. A. Jares-Erijman, T. M. Jovin, Nat. Biotechnol. 2003, 21, 1387.

[14] A. Margineanu, J. J. Chan, D. J. Kelly, S. C. Warren, D. Flatters, S. Kumar, M. Katan, C. W. Dunsby, P. M. W. French, Sci. Rep. 2016, 6, 33621.

[15] T. W. J. Gadella, T. M. Jovin, R. M. Clegg, Biophys. Chem. 1993, 48, 221.

[16] W. Becker, J. Microsc. 2012, 247, 119.

[17] L. Marcu, Ann. Biomed. Eng. 2012, 40, 304.

[18] S. L. Jacques, Phys. Med. Biol. 2013, 58, R37.

[19] M. A. Digman, V. R. Caiolfa, M. Zamai, E. Gratton, Biophys. J. 2008, 94, L14.

[20] F. Fereidouni, A. Esposito, G. A. Blab, H. C. Gerritsen, J. Microsc. 2011, 244, 248.

[21] N. G. James, J. A. Ross, M. Stefl, D. M. Jameson, Anal. Biochem. 2011, 410, 70.

[22] C. Stringari, A. Cinquin, O. Cinquin, M. A. Digman, P. J. Donovan, E. Gratton, P. Natl. Acad. Sci. USA 2011, 108, 13582.

[23] R. A. Colyer, O. H. W. Siegmund, A. S. Tremsin, J. V. Vallerga, S. Weiss, X. Michalet, J. Biomed. Opt. 2012, 17, 016008.

[24] E. Hinde, M. A. Digman, K. M. Hahn, E. Gratton, P. Natl. Acad. Sci. USA 2013, 110, 135.

[25] D. M. Jameson, C. M. Vetromile, N. G. James, Methods 2013, 59, 278.

[26] F. Fereidouni, A. N. Bader, A. Colonna, H. C. Gerritsen, J. Biophotonics 2014, 7, 589.

[27] H. Szmacinski, V. Toshchakov, J. R. Lakowicz, J. Biomed. Opt. 2014, 19, 046017.

[28] F. Cutrale, V. Trivedi, L. A. Trinh, C. L. Chiu, J. M. Choi, M. S. Artiga, S. E. Fraser, Nat. Methods 2017, 14, 149.

[29] F. Fereidouni, D. Gorpas, D. Ma, H. Fatakdawala, L. Marcu, Methods Appl. Fluoresc. 2017, 5, 035003.

[30] S. Osseiran, E. M. Roider, H. Q. Wang, Y. Suita, M. Murphy, D. E. Fisher, C. L. Evans, J. Biomed. Opt. 2017, 22, 125004.

[31] F. Radaelli, L. D’Alfonso, M. Collini, F. Mingozzi, L. Marongiu, F. Granucci, I. Zanoni, G. Chirico, L. Sironi, Sci. Rep. 2017, 7, 17468.

[32] C. A. Gomez, J. Sutin, W. C. Wu, B. Y. Fu, H. Uhlirova, A. Devor, D. A. Boas, S. Sakadzic, M. A. Yaseen, Plos One 2018, 13, e0194578.

[33] S. Sameni, L. Malacrida, Z. Q. Tan, M. A. Digman, Sci. Rep. 2018, 8, 734.

[34] T. R. Daniels, T. Delgado, J. A. Rodriguez, G. Helguera, M. L. Penichet, Clin Immunol 2006, 121, 144.

[35] T. R. Daniels, E. Bernabeu, J. A. Rodriguez, S. Patel, M. Kozman, D. A. Chiappetta, E. Holler, J. Y. Ljubimova, G. Helguera, M. L. Penichet, Biochim. Biophys. Acta 2012, 1820, 291.

[36] A. Berczi, K. Barabas, J. A. Sizensky, W. P. Faulk, Arch. Biochem. Biophys. 1993, 300, 356.

[37] M. Weaver, D. W. Laske, J. Neuro-Oncol. 2003, 65, 3.

[38] H. Wallrabe, G. Bonamy, A. Periasamy, M. Barroso, Mol. Biol. Cell 2007, 18, 2226.

[39] R. Talati, A. Vanderpoel, A. Eladdadi, K. Anderson, K. Abe, M. Barroso, Methods 2014, 66, 139.

[40] K. Abe, L. L. Zhao, A. Periasamy, X. Intes, M. Barroso, Plos One 2013, 8, e80269.

[41] A. Rudkouskaya, N. Sinsuebphon, J. Ward, K. Tubbesing, X. Intes, M. Barroso, J. Control. Release 2018, 286.

[42] W. J. Akers, M. Y. Berezin, H. Lee, S. Achilefu, J. Biomed. Opt. 2008, 13, 054042.

[43] W. Becker, Advanced time-correlated single photon couting techniques, Springer,Berlin, 2005.

[44] B. Heeg, Meas. Sci. Technol. 2014, 25, 105201.

[45] A. Leray, S. Padilla-Parra, J. Roul, L. Heliot, M. Tramier, Plos One 2013, 8, e69335.

[46] S. Padilla-Parra, N. Auduge, M. Coppey-Moisan, M. Tramier, 2011, 3, 63.

[47] D. M. Jameson, E. Gratton, R. D. Hall, Appl. Spectrosc. Rev. 1984, 20, 55.

[48] G. I. Redford, R. M. Clegg, J. Fluoresc. 2005, 15, 805.

[49] G. Weber, J. Phys. Chem. 1981, 85, 949.

[50] A. Xiao, A. E. Gibbons, K. E. Luker, G. D. Luker, Tomography 2015, 1, 115.

[51] A. H. A. Clayton, Q. S. Hanley, P. J. Verveer, J. Microsc. 2004, 213, 1.

[52] R. A. Colyer, C. Lee, E. Gratton, Microsc. Res. Techniq. 2008, 71, 201.

[53] E. Hinde, M. A. Digman, C. Welch, K. M. Hahn, E. Gratton, Microsc. Res. Techniq. 2012, 75, 271.

[54] T. Hato, S. Winfree, R. Day, R. M. Sandoval, B. A. Molitoris, M. C. Yoder, R. C. Wiggins, Y. Zheng, K. W. Dunn, P. C. Dagher, J. Am. Soc. Nephrol. 2017, 28, 2420.

[55] F. J. Vergeldt, A. Prusova, F. Fereidouni, H. van Amerongen, H. van As, T. W. J. Scheenen, A. N. Bader, Sci. Rep. 2017, 7, 861.

[56] L. L. Zhao, K. Abe, M. Barroso, X. Intes, Opt. Lett. 2013, 38, 3976.

[57] V. Venugopal, J. Chen, X. Intes, Biomed. Opt. Express 2010, 1, 143.

[58] L. L. Zhao, K. Abe, S. Rajoria, Q. Pian, M. Barroso, X. Intes, Biomed. Opt. Express 2014, 5, 944.

[59] S.-J. Chen, N. Sinsuebphon, X. Intes, Photonics 2015, 2, 1027.

[60] A. Furstenberg, Chimia 2017, 71, 26.

[61] S. F. Lee, Q. Verolet, A. Furstenberg, Angew. Chem. Int. Ed. 2013, 52, 8948.

[62] L. A. A. de Jong, D. R. A. Uges, J. P. Franke, R. Bischoff, J. Chromatogr. B 2005, 829, 1.

[63] A. D. Goddard, A. Watts, Biophys. Rev. 2012, 4, 291.

[64] Z. M. Qian, H. Y. Li, H. Z. Sun, K. Ho, Pharmacol. Rev. 2002, 54, 561.

[65] J. F. Cormier, M. Fortin, J. Frechette, I. Noiseux, M. L. Vernon, W. Long, Proc. SPIE 2005, 5702, 123.

[66] C. L. Hutchinson, T. L. Troy, E. M. Sevick-Muraca, Appl. Optics 1996, 35, 2325.

[67] V. Venugopal, J. Chen, M. Barroso, X. Intes, Biomed. Opt. Express 2012, 3, 3161.

[68] V. Venugopal, J. Chen, F. Lesage, X. Intes, Opt. Lett. 2010, 35, 3189.

[69] C. Bruschini, H. Homulle, E. Charbon, Proc. SPIE 2017, 10069, 10069S.

[70] D. G. Myszka, X. He, M. Dembo, T. A. Morton, B. Goldstein, Biophys. J. 1998, 75, 583.

[71] A. P. West, M. J. Bennett, V. M. Sellers, N. C. Andrews, C. A. Enns, P. J. Bjorkman, J. Biol. Chem. 2000, 275, 38135.

[72] T. Inoue, P. G. Cavanaugh, P. A. Steck, N. Brunner, G. L. Nicolson, J. Cell Physiol. 1993, 156, 212.

